# The Influence of Blood Collection Tubes in Biomarkers Screening by Mass Spectrometry

**DOI:** 10.1101/753111

**Authors:** Siyuan Zhang, Zixuan Zhao, Wenjing Duan, Zhaoxin li, Zhuhui Nan, Hanzhi Du, Mengchang Wang, Juan Yang, Chen Huang

## Abstract

**Background:** Mass spectrometry (MS) is one of the rapidly developing bio-analytical techniques in recent years and have found many biomarkers of variety of disease. Whereas pre-analytical process is one of most crucial procedure which would significantly influence the results of biomarkers screening. In the current study, we conducted a pilot analysis of serum to determine the effects of blood collection tubes in biomarkers screening.

**Methods:** Magnetic bead separation and matrix-assisted laser desorption ionization time-of-flight mass spectrometry were used for qualitative analysis of healthy control and serum cancer patients. A total of 24 serum samples were analyzed in this study, of which were collected from patients or healthy control using non-additive tubes or coagulant activator tubes respectively. ClinProTools were used to compare the difference among the different groups.

**Results:** The results demonstrated that no matter for patients or normal people, the serum protein profile changed significantly when using coagulant tubes. We also found that the effect of coagulant on serum protein of patients was smaller than that of control group. There were significant differences among 27 peaks which were obtained in the control group and the control coagulant group. However, between patient group and patient coagulant group, only 1 differential peak were obtained. Coagulant changed the protein expression difference in the original serum, and the difference expanded, narrowed even reversed, most of which are small polypeptides (Mass<3000 Da), which significantly changed the results of biomarkers screening. The results showed that 19 potential biomarkers could be found with non-additive tubes and 16 potential biomarkers could be found with coagulate activator tubes, among which only 6 were the same.

**Conclusions:** The choice of blood collection tube significantly influence the results of biomarkers screening by MS.

## Introduction

Tumor is a complex disease caused by multiple factors and regulated by many genes^1,2^. Genes are expressed in complex ways, while proteins are the executors of cell function and can dynamically reflect biological systems. Therefore, the formation of tumor cells must be related to proteome changes. Blood is a convenient source of biomarkers which is readily available, and it is immersed in most tissues of the body. Therefore, it probably contains cell-derived proteins and peptides that provide information about various biological processes. Protein biomarkers, which provide molecular signatures of certain diseases, can be used for diagnosis (early detection), prognosis (staging detection), determining efficacy (translational research), monitoring of toxicity, and screening of drug targets^3,4^.

Low molecular weight (LMW, molecular weight⩽30kDa) proteins /peptidome, mostly in the serum^5^, contain an unexplored archive of histological information and expected for useful biomarkers screening in early detection^6^. Mass spectrometry (MS) is one of the rapidly developing bio-analytical techniques in recent years, which has been used to identify and validate protein biomarkers for clinical application^7^. MS-based proteomics can detect the whole protein system, providing a useful window for understanding a series of biological processes and allowing the identification of differentially expressed proteins between samples^8^. As a method of detecting biological active molecules, MS has many advantages^9^. In addition, the characteristics of mass spectrometry can be summarized as sensitivity, rapidity, specificity and high throughput, which fully demonstrates that MS is superior to other methods.

Though such multiplex profiling and modeling methods have found a lot of biomarkers of variety of diseases, the concerns about the quality control have been raised^10,11^. Several novel algorithms and software have been invented to solve the problem of reproducibility and reliability of this promising technology^12-14^. Furthermore, pre-analytical serum sample procedures is another important concerns, only a few studies have focused on the effects of pre-analytical conditions and put forward some approach^15-17^, other potential influence factors included storage temperature^18^ and time^19,20^, time intervals between venipuncture and serum preparation^21^, and the type of blood collection tubes^22-24^. There are variety types of Blood collection tube wildly used in clinical field, while, totally speaking, coagulant tubes or anticoagulant tubes are used for collecting serum or plasma, respectively^25^. Since most of the LMW proteins are in the serum, the non-additive tube (nature coagulation) and coagulant tube are two main collection tubes. Previous study^23^ have determined the results between the two types were not the same. However, previous studies only tested the influence of pre-analytical process in healthy controls and neglected the influence of patients and comparison between them. Thus, these previous results are questionable that whether such different pre-analytical process would influence the results of potential biomarker screening.

In the current study we determined whether the type of blood collection tubes influence the results of potential biomarker finding. We used matrix-assisted laser desorption ionization-time of flight mass spectrometry (MALDI-TOF MS) and ClinProTools software to compare the difference of serum peptidome profiles across samples from 6 healthy controls and 6 patients whose blood were collected directly into two different types of blood collection tubes (coagulant activator tube and tube without any additive). Understanding the influence of pre-analytical procedures to potential biomarker finding will give us a clear idea when formulate and conduct pre-analytical procedures.

## Materials and methods

### Patients and Sample Preparation

The study protocol was approved by the ethics committee and the human research review committee of Xi’an Jiao tong University. A total of 6 serum samples of serum cancer patients (3 men and 3 women) including 2 multiple myeloma patients, 2 non-Hodgkin lymphoma patients, 1 acute B lymphoblastic leukemia and 1 acute leukemia, were obtained from the First Affiliated Hospital of Xi’an Jiao tong University. The range for patients was from 54 to 73 years with an average age of 64.7. All the patients were recently diagnosed. Serum samples from 6 healthy controls, consisting of 2 men and 4 women ranging in age from 19 to 20 years old, were obtained from recruited healthy donors. All blood samples were collected while the patients or healthy controls were seated and fasting. The samples were collected in coagulant activator and non-additive tube, then centrifuged at 3,000 rpm for 20 min at 4°C. The serum samples were distributed into 500 μL aliquots and stored at −80°C until use.

### WCX Fractionation and MALDI-TOF MS

WCX fractionation and MALDI-TOF mass spectrometry combined. This experiment used a magnetic bead based weak cation exchange chromatography (MB-WCX) to separate the sample and used a Klingput purification reagent. The MB-WCX purifier used the Bruker magnet to execute according to the manufacturer’s agreement. Using MB-WCX and magnetic pickers, magnets reduced 5 μL serum sample dilution 10 μL binding in a standard PCR tube, and added to 10 μL MB-WCX beads. After fully stirring, incubated the sample at room temperature for 5 min, then placed the test tube in the magnetic separator, collected the beads on the test tube wall, and clarified the upper liquid (1 min). Then removed the liquid and lower the magnet again. After a series of related operations, we screened the peptide fraction from the magnetic beads of 5 μL elution solution and 5 μL stable buffer. After the step-by-step application example and the MB-WCX separation, we screened the peptide fraction from the magnetic bead 5μL elution solution and the 5μL stable buffer. The screening of peptides was found on the surface of MALDI AnchoChip 1 μ L alpha-cyano-4-hydroxy cinnamic acid (force Daltonics) in 50 % acetonitrile, and 0.5 % trifluoroacetic acid MALDI AnchaorChip tripled. In order to evaluate reproducibility, each sample was marked in triplicate.

### MS data analysis

MS data analyzed of all processed samples, and then immediately data processing with Clinprot software, air-dry target immediately with the calibrated Autoflex III MALDI-TOF MS (Bruker), Flex-Control software (version 3.0 and the associated measurement protocols. The matrix inhibits up to 700da, and the mixture which the mass range 800-10000Da is calibrated with polypeptide and protein standards for quality calibration. The blind method was used for all measurements, including a mixture of serum in the patient and control group. Data analyse used Flex analysis software (version on 3.0; Use Bruker). ClinProTools software (version 2.2) was software for peptide pattern recognition. The program used a standard data processing model, including spectral preprocessing, peak extraction and peak calculation operations.

## Results

In our study, we analyzed the serum proteomic profiles of all the four groups, including the patient group, control group, patient coagulant group and control coagulant group. We evaluated the changes at the peptidome level in the serum samples of the patient group, control group in comparison to their respective coagulant group, and analyzed differences between the patient group, patient coagulant group, and their respective controls and eventually analyzed the differences among all the four groups. By analyzing the spectra (screened from different groups) using ClinProTools software 2.2, we were able to confirm that the type of serum separate tubes significantly influences the results of biomarkers screening by mass spectrometry.

### Serum profiles of the four groups and assay reproducibility

In this study, all the samples were prefractionated by MB-WCX magnetic beads and detected by MALDI-TOF MS respectively. It also showed that all the groups had proteomic profiles from 800 to 10,000 Da in which a significant number of differentially expressed peaks can be easily detected. We had also performed the reproducibility and stability of the mass spectra of this study. All the samples were in triplicate for both MB-WCX and MALDI-TOF MS purification and showed closely reproducible peaks (Figure 1).

**Figure 1.**
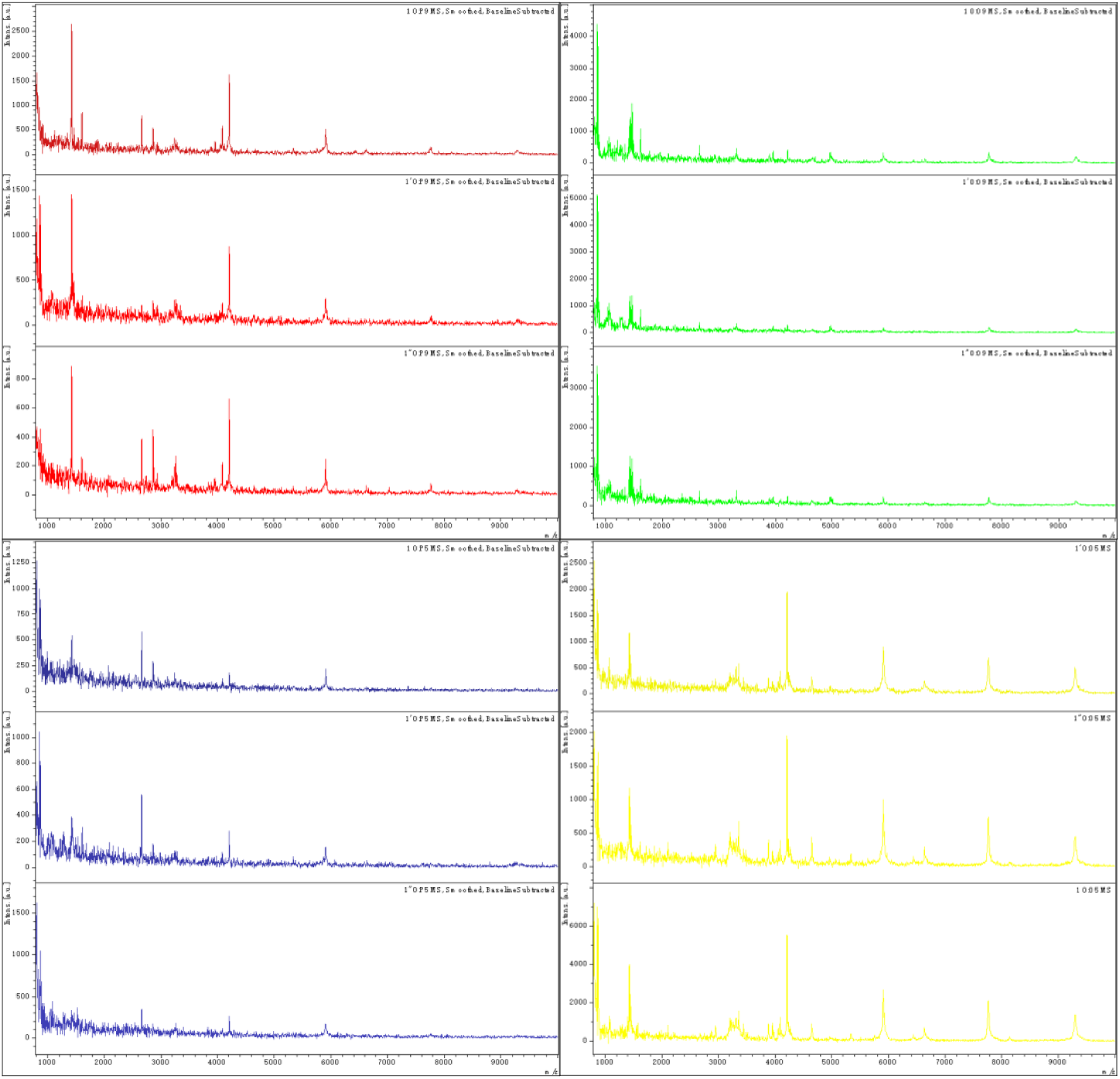
Comparative profiling of serum peptides among patient group (red), control group (green), patient coagulant group (blue) and control coagulant group (yellow). Representative mass spectra of the samples (three spectra per sample) from different groups respectively (differ in colors) in the mass range of 800–10,000 Da, showing low variability between replicates of each sample.

### Coagulant significantly affect serum protein spectrum

We found that there were some different peaks among patient group, control group, patient coagulant group and control coagulant group by ClinProTools. A total of 61 peaks of which there were significant differences among 27 peaks were obtained in the control group and the control coagulant group (Table 2). In the 27 peaks, 21 peaks were up-regulated (Figure2 D) and 6 peaks were down-regulated (Figure2 E). Among them, the fold changes of 14 peak polypeptides were more than 1.5 times in which the largest difference was 2.66 times and the molecular mass of the polypeptides was 5927.63. In addition, the remaining peaks were peak 2 (Mass:4237.14), peak 3 (Mass:4256.45), peak 4 (Mass:3365.31), peak 6 (Mass:4272.8), peak 7 (Mass:5871.62), peak 8 (Mass:5911.94), peak 10 (Mass:4215.61), peak 11 (Mass:3236.73), peak 13 (Mass:3212.5), peak 15 (Mass:6638.57), peak 19 (Mass:9302.34), peak 20 (Mass:7774.02) and peak 21 (Mass:810.18). Four peptides showed the difference of less than 0.67, and the peaks included peak 5 (Mass: 4970.81), peak 9 (Mass: 4989.4), peak 12 (Mass: 1620.29) and peak 18 (Mass: 1468.98). (Table 2) Between patient group and patient coagulant group, only 1 differential down-regulated peaks was obtained (Table 1, Figure 3D). It’s worth mentioning that there was an overlap between the bivariate and three-dimensional diagrams of the patient group and the patient coagulant group (Figure 3B C). Therefore, the comparison and analysis of the two groups of experiments showed that the effect of coagulant on serum protein of patients was smaller than that of control group.

**Table 1.** Peaks showing significant differences in abundance across samples from patient coagulant and patient.

| Peak | Mass | P value | patient<br>coagulant | patient | fold change |
| --- | --- | --- | --- | --- | --- |
| 1 | 6638.04 | 0.0159 | $0.73 \pm 0.13$ | $1.02 \pm 0.28$ | 0.72 |

**Table 2.** Peaks showing significant differences in abundance across samples from control coagulant and control.

| Peak | Mass | P value | control<br>coagulant | control | fold change |
| --- | --- | --- | --- | --- | --- |
| 1 | 5927.63 | < 0.000001 | $3.25 \pm 0.93$ | $1.22 \pm 0.37$ | 2.66* |
| 2 | 4237.14 | < 0.000001 | $2.57 \pm 0.59$ | $1.4 \pm 0.31$ | 1.84* |
| 3 | 4256.45 | 0.00000575 | $1.83 \pm 0.34$ | $1.2 \pm 0.28$ | 1.53* |
| 4 | 3365.31 | 0.00000656 | $3.05 \pm 0.98$ | $1.45 \pm 0.3$ | 2.10* |
| 5 | 4970.81 | 0.0000114 | $0.81 \pm 0.22$ | $1.8 \pm 0.58$ | 0.45* |
| 6 | 4272.8 | 0.0000153 | $1.89 \pm 0.48$ | $1.11 \pm 0.33$ | 1.70* |
| 7 | 5871.62 | 0.0000322 | $1.44 \pm 0.37$ | $0.9 \pm 0.2$ | 1.60* |
| 8 | 5911.94 | 0.0000356 | $5.06 \pm 2.22$ | $1.97 \pm 0.92$ | 2.57* |
| 9 | 4989.4 | 0.0000356 | $0.77 \pm 0.17$ | $1.39 \pm 0.41$ | 0.55* |
| 10 | 4215.61 | 0.000062 | $8.28 \pm 3.68$ | $3.35 \pm 1.65$ | 2.47* |
| 11 | 3236.73 | 0.00017 | $2.19 \pm 0.58$ | $1.45 \pm 0.32$ | 1.51* |
| 12 | 1620.29 | 0.000199 | $2.86 \pm 0.92$ | $5.5 \pm 2.04$ | 0.52* |
| 13 | 3212.5 | 0.000263 | $2.52 \pm 0.74$ | $1.65 \pm 0.33$ | 1.53* |
| 14 | 5831.2 | 0.00042 | $0.95 \pm 0.2$ | $0.7 \pm 0.14$ | 1.36 |
| 15 | 6638.57 | 0.000443 | $1.3 \pm 0.53$ | $0.72 \pm 0.15$ | 1.81* |
| 16 | 3283.36 | 0.000476 | $2.28 \pm 0.46$ | $1.74 \pm 0.32$ | 1.31 |
| 17 | 3340.88 | 0.000751 | $2.25 \pm 0.49$ | $1.66 \pm 0.38$ | 1.36 |
| 18 | 1468.98 | 0.00129 | $4.59 \pm 1.26$ | $7.67 \pm 2.9$ | 0.60* |
| 19 | 9302.34 | 0.00164 | $2.19 \pm 0.92$ | $1.23 \pm 0.59$ | 1.78* |
| 20 | 7774.02 | 0.00189 | $3.38 \pm 1.38$ | $1.9 \pm 1.02$ | 1.78* |
| 21 | 810.18 | 0.0029 | $18.38 \pm 9.17$ | $10.25 \pm 2.87$ | 1.79* |
| 22 | 818.37 | 0.00926 | $7.93 \pm 2.04$ | $6.16 \pm 1.39$ | 1.29 |
| 23 | 834.9 | 0.00948 | $8.72 \pm 2.76$ | $6.49 \pm 1.45$ | 1.34 |
| 24 | 2664.58 | 0.00949 | $3.02 \pm 1.47$ | $5.49 \pm 2.96$ | 0.55 |
| 25 | 3957.94 | 0.014 | $1.54 \pm 0.59$ | $2.07 \pm 0.52$ | 0.74 |
| 26 | 5343.35 | 0.0315 | $1.19 \pm 0.39$ | $0.88 \pm 0.33$ | 1.35 |
| 27 | 4651.16 | 0.0497 | $1.89 \pm 0.79$ | $1.41 \pm 0.38$ | 1.34 |

**Figure 2.**
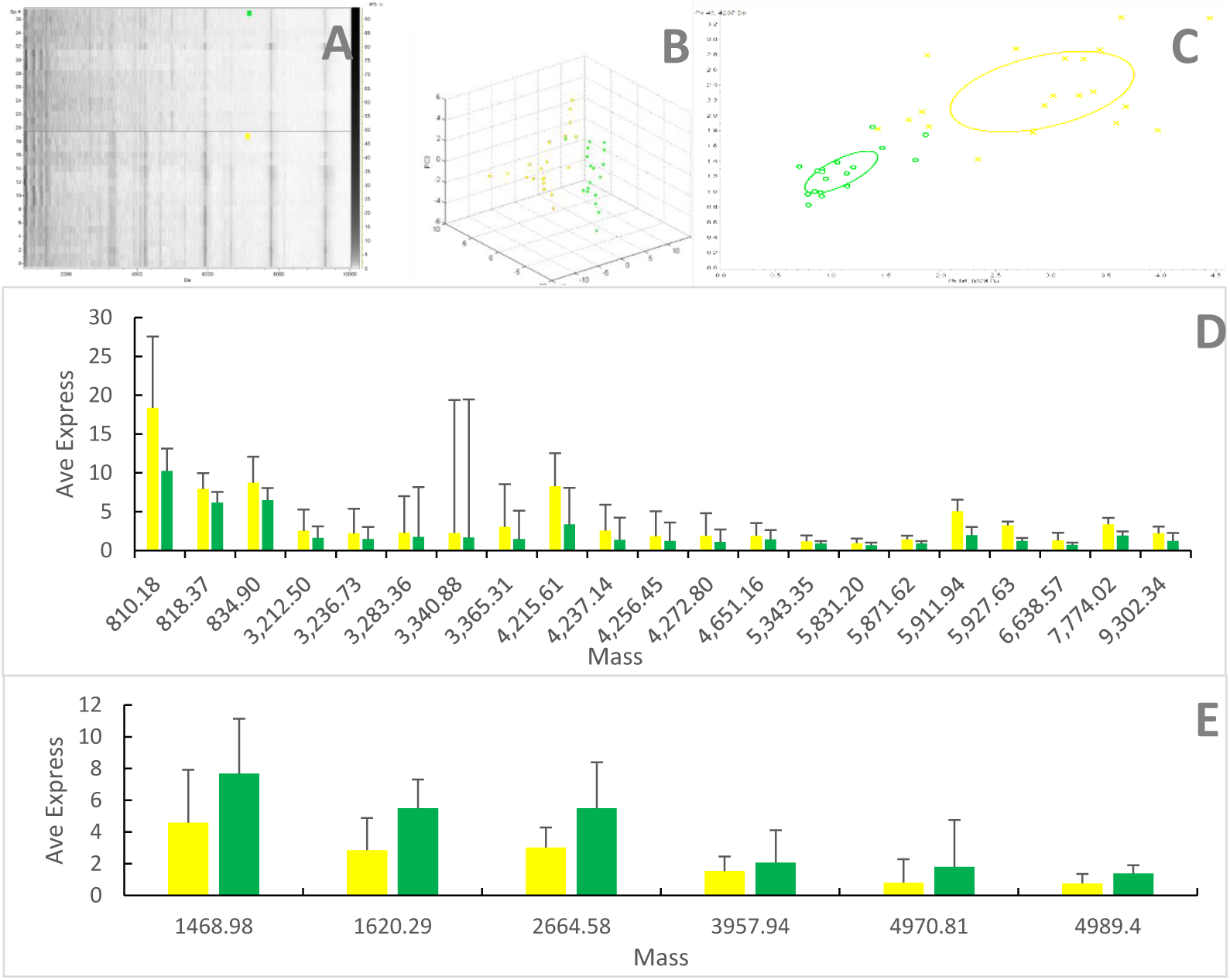
Comparison of serum protein in control group (green) and control coagulant group (yellow. (A) The molecular weight of Control group (green) and control coagulant group (yellow) of expressed peptides ranged from 800 to 10000 Da. Bivariate plot (C) and 3D plot (B) of control group (green) and control coagulant group (yellow) in the principal component analysis. Polypeptide expression in control group (green) and control coagulant group (yellow) was up-regulated (D), or down-regulated (E).

**Figure 3.**
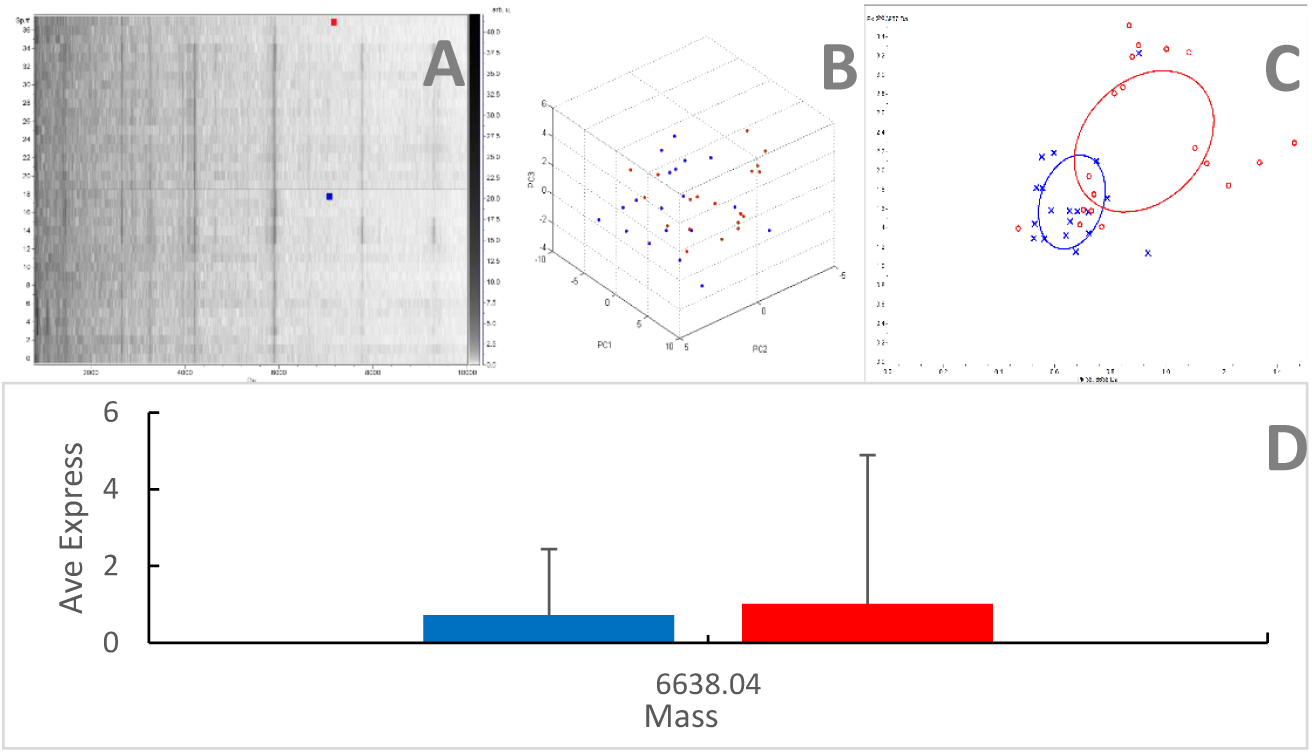
Comparison of serum protein in patient group (red) and patient coagulant group (blue). (A)The molecular weight of patient group (red) and patient coagulant group (blue) of expressed peptides ranged from 800 to 10000 Da. Bivariate plot (C) and 3D plot (B) of patient group (red) and patient coagulant group (blue) in the principal component analysis. Polypeptide expression in patients with non-additive tubes and patients with coagulant activators was down-regulated (D).

### The significantly changed peaks between patient and control sera collected by coagulant or non-additive tubes

The results from the comparison of the samples of control group and patient group showed that the potential serum markers collected by the coagulant activator or non-additive tube were not exactly the same. A large number of potential serum markers were found in the serum of patient group with control group and patient coagulant group with control coagulant group collected with non-additive tube and the coagulant activator tube.

There is basically no overlap between the peak of patient group and control group (Figure 4C and Figure 5C). Compared with control group, there were 12 up-regulated peaks in the samples collected by patients with non-additive tube (Figure 5D), and there were 13 up-regulated peaks in the samples collected by patients with the coagulant activator (Figure 4D), but none of them were the same. Compared with the samples of control group, there were 14 down-regulated peaks in the samples collected by the non-additive tube (Figure 5E), but 23 down-regulated peaks in the serum collected by the coagulant activator (Figure 4E), only one of which was the same. This showed that the use of the coagulant activator to collect blood samples had a significant impact on the results.

**Figure 4.**
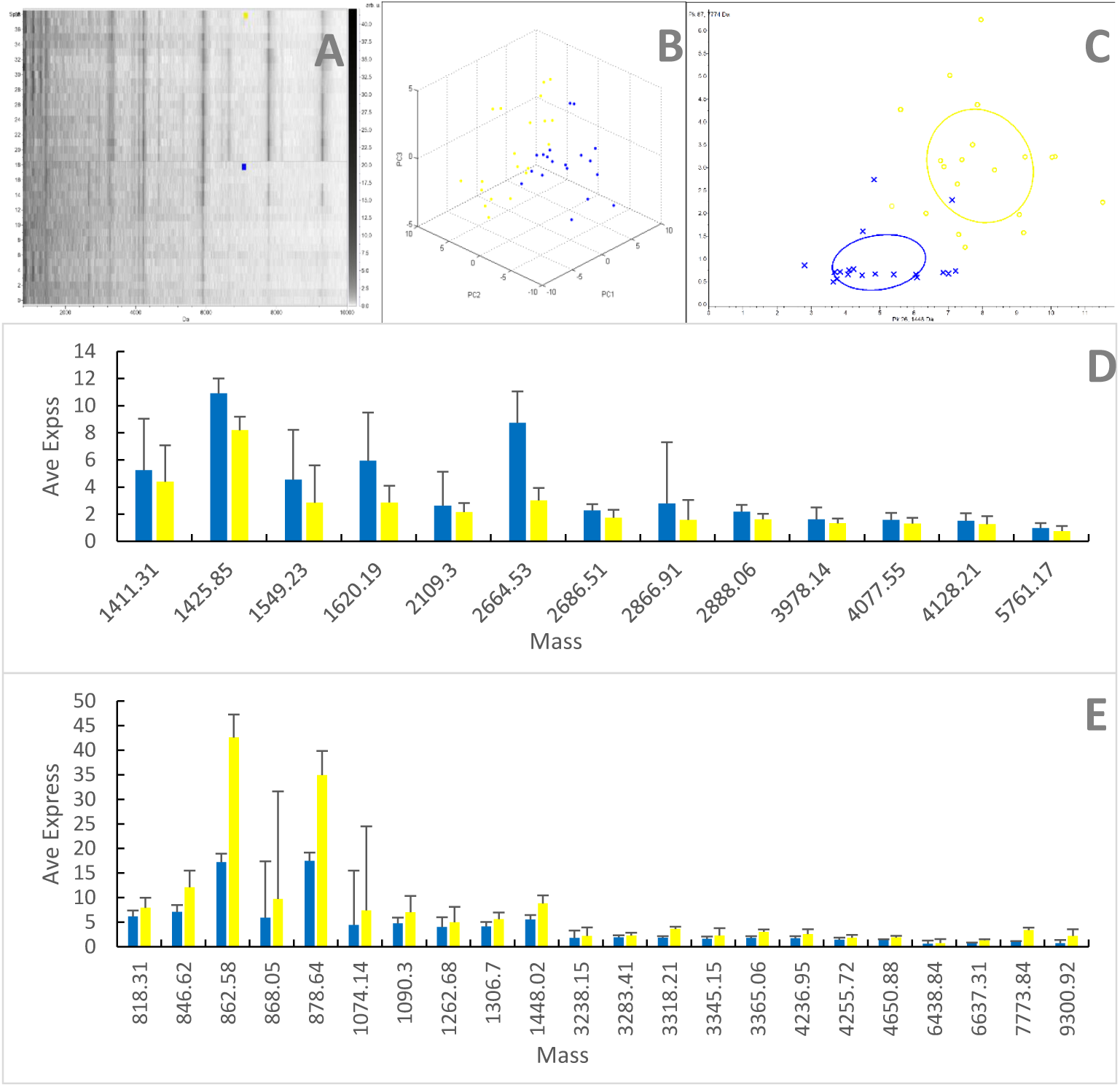
Comparison of serum protein collected by coagulant tubes in patient group (blue) and control group (yellow). (A)Gel view of mass spectra of patient coagulant group and control coagulant group in the mass range from 800 to 10,000 Da. (B) Three-dimensional plot of patient coagulant group (blue) and control coagulant group (yellow). (C)Bivariate plot of the two most differentially expressed protein peaks of patient coagulant group (blue) and control coagulant group (yellow). (D) Collected by patient coagulant group (blue) and control coagulant group (yellow) with higher expression in serum protein expression differences between the pillars of the mean and standard deviation of the peak draw diagrams. (E) Collected by patient coagulant group (blue) and control coagulant group (yellow) with lower expression of serum protein expression differences between the pillars of the mean and standard deviation of the peak draw diagrams.

**Figure 5.**
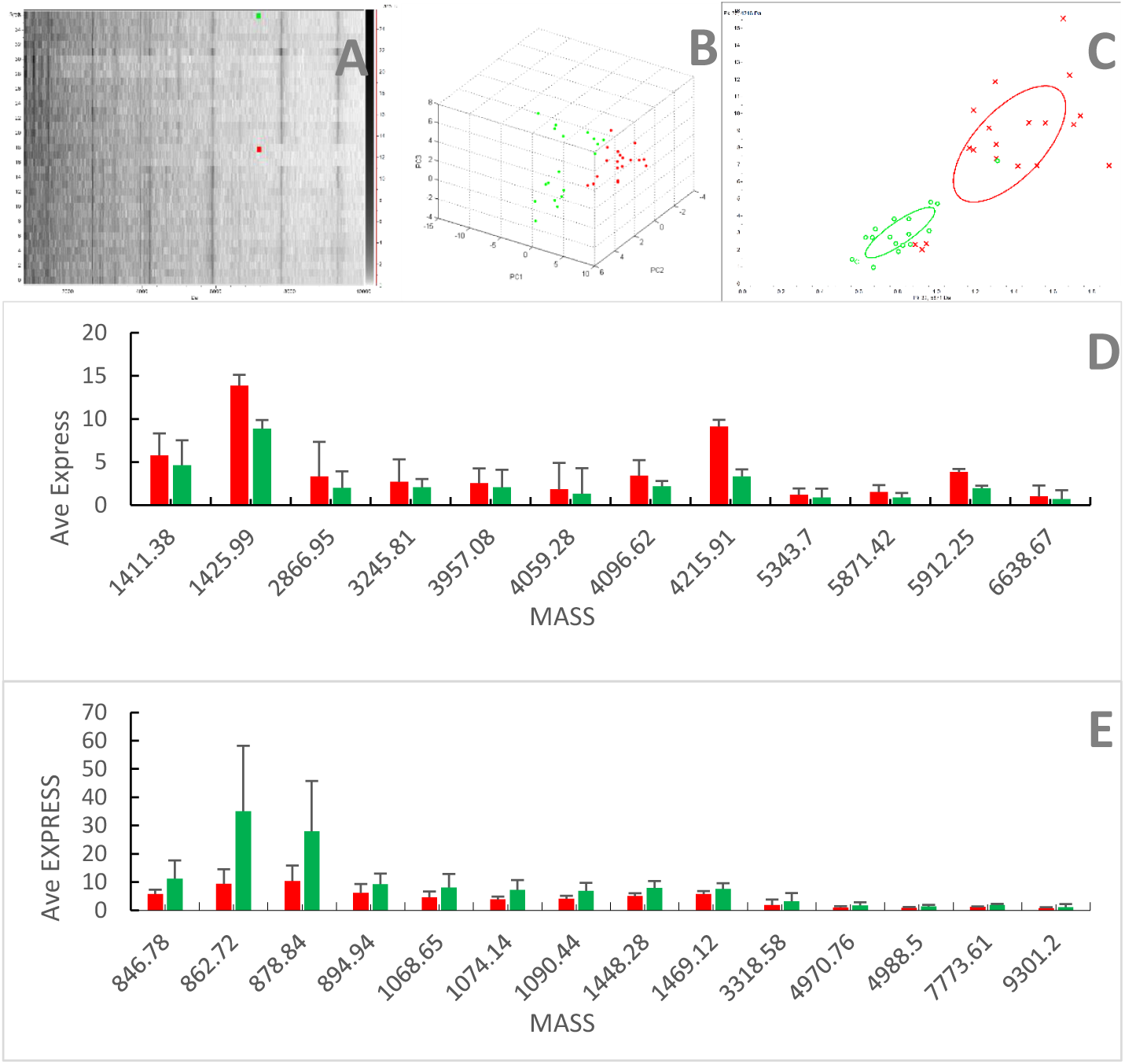
Comparison of serum protein collected by non-additive tubes in patient group (red) and control group (green). (A) Gel view of mass spectra of patient group and control group in the mass range from 800 to 10,000 Da. (B) Three-dimensional plot of patient group (red) and control group (green). (C) Bivariate plot of the two most differentially expressed protein peaks of patient group (red) and control group (green). (D) Collected by non-additive tube (red) and healthy controls (green) in patients with higher expression in serum protein expression differences between the pillars of the mean and standard deviation of the peak draw diagrams. (E) Collected by non-additive tube (red) and healthy controls (green) in patients with lower expression of serum protein expression differences between the pillars of the mean and standard deviation of the peak draw diagrams.

In addition, we found some peaks with significant differences. This showed that the use of the coagulant activator to collect blood samples had a significant impact on the results of biomarker screening. In addition, we found some peaks with significant differences. In the table 3, according to statistics, there were 19 peaks with multiple difference greater than 1.5 or less than or equal to 0.67, in which 6 peaks were significantly up-regulated and 13 were significantly down-regulated. The up-regulated peaks are peak 1 (Mass: 5871.42), peak 2(Mass: 4215.91), peak 4(Mass: 1425.99), peak 5(Mass: 5912.25), peak 17(Mass: 4096.62), peak 21(Mass: 2866.95). And there were 13 significantly lower peaks, peak 3(Mass: 1448.28), peak 6(Mass: 3318.58), peak 8(Mass: 4970.76), peak 9(Mass: 1074.14), peak 10(Mass: 1090.44), peak 11(Mass: 4988.5), peak 12(Mass: 862.72), peak 14(Mass: 1068.65), peak 15(Mass: 878.84), peak 16(Mass: 846.78), peak 20(Mass: 9301.2), peak 22(Mass: 7773.61), peak 23(Mass: 894.94). In the table 4, according to statistics, there were 16 peaks with multiple difference greater than 1.5 or less than or equal to 0.67, in which four peaks were significantly up-regulated and 12 were significantly down-regulated. The up-regulated peaks were peak 4 (Mass: 2866.91), peak 7 (Mass: 1620.19), peak 9 (Mass: 2664.53) and peak 23 (Mass: 1549.23). Peak 9 had the largest upward revision. And there are 12 significantly lower peaks, peak 1 (Mass: 1448.02), the peak 2 (Mass: 7773.84), the peak 3 (Mass: 9300.92), peak 5 (Mass: 4236.95), peak 6 (Mass: 3365.06), peak 10 (Mass: 3318.21), peak 11 (Mass: 862.58), peak 13(Mass: 6637.13), the peak 14 (Mass: 846.62), the peak 15 (Mass: 878.64), peak 18 (Mass: 1074.14), the peak 21 (Mass:868.05). And peak 2 had the largest downward range.

**Table 3.** Peaks showing significant differences in abundance across samples from patient and control.

| Peak | Mass | P value | patient | control | fold change |
| --- | --- | --- | --- | --- | --- |
| 1 | 5871.42 | 0.00000192 | $1.53 \pm 0.32$ | $0.9 \pm 0.2$ | 1.70* |
| 2 | 4215.91 | 0.0000447 | $9.13 \pm 3.79$ | $3.35 \pm 1.65$ | 2.73* |
| 3 | 1448.28 | 0.0000753 | $5.05 \pm 1.03$ | $7.98 \pm 1.95$ | 0.63* |
| 4 | 1425.99 | 0.000228 | $13.89 \pm 4.01$ | $8.88 \pm 1.91$ | 1.56* |
| 5 | 5912.25 | 0.000228 | $3.85 \pm 1.44$ | $1.97 \pm 0.92$ | 1.95* |
| 6 | 3318.58 | 0.000228 | $1.86 \pm 0.4$ | $3.23 \pm 1.06$ | 0.58* |
| 7 | 4059.28 | 0.000228 | $1.84 \pm 0.35$ | $1.34 \pm 0.3$ | 1.37 |
| 8 | 4970.76 | 0.00035 | $1.08 \pm 0.27$ | $1.8 \pm 0.58$ | 0.60* |
| 9 | 1074.14 | 0.000365 | $3.85 \pm 1.05$ | $7.26 \pm 2.84$ | 0.53* |
| 10 | 1090.44 | 0.000576 | $4.13 \pm 1.02$ | $6.88 \pm 2.4$ | 0.60* |
| 11 | 4988.5 | 0.000576 | $0.92 \pm 0.2$ | $1.39 \pm 0.41$ | 0.66* |
| 12 | 862.72 | 0.000598 | $9.41 \pm 5.14$ | $35.13 \pm 23.11$ | 0.27* |
| 13 | 6638.67 | 0.00096 | $1.02 \pm 0.28$ | $0.72 \pm 0.15$ | 1.42 |
| 14 | 1068.65 | 0.0014 | $4.65 \pm 1.06$ | $8.15 \pm 3.46$ | 0.57* |
| 15 | 878.84 | 0.00163 | $10.42 \pm 5.47$ | $28 \pm 17.8$ | 0.37* |
| 16 | 846.78 | 0.00566 | $5.86 \pm 1.45$ | $11.27 \pm 6.43$ | 0.52* |
| 17 | 4096.62 | 0.00724 | $3.42 \pm 1.28$ | $2.21 \pm 1.03$ | 1.55* |
| 18 | 1411.38 | 0.00762 | $5.8 \pm 1.24$ | $4.66 \pm 1$ | 1.24 |
| 19 | 5343.7 | 0.00762 | $1.21 \pm 0.3$ | $0.89 \pm 0.33$ | 1.36 |
| 20 | 9301.2 | 0.01 | $0.75 \pm 0.33$ | $1.2 \pm 0.56$ | 0.63* |
| 21 | 2866.95 | 0.0107 | $3.35 \pm 1.81$ | $2.02 \pm 0.6$ | 1.66* |
| 22 | 7773.61 | 0.0213 | $1.21 \pm 0.38$ | $1.9 \pm 1.02$ | 0.64* |
| 23 | 894.94 | 0.0225 | $6.16 \pm 2.03$ | $9.34 \pm 4.71$ | 0.66* |
| 24 | 3245.81 | 0.037 | $2.71 \pm 0.78$ | $2.09 \pm 0.82$ | 1.30 |
| 25 | 1469.12 | 0.0411 | $5.79 \pm 1.98$ | $7.67 \pm 2.9$ | 0.75 |
| 26 | 3957.08 | 0.0459 | $2.57 \pm 0.81$ | $2.07 \pm 0.52$ | 1.24 |

**Table 4.** Peaks showing significant differences in abundance across samples from patient coagulant and control coagulant.

| Peak | Mass | P value | patient<br>coagulant | control<br>coagulant | fold change |
| --- | --- | --- | --- | --- | --- |
| 1 | 1448.02 | 0.00000825 | 5.53±1.51 | 8.81±1.73 | 0.63* |
| 2 | 7773.84 | 0.00000825 | 1.03±0.68 | 3.38±1.38 | 0.30* |
| 3 | 9300.92 | 0.0000126 | 0.71±0.41 | 2.19±0.92 | 0.32* |
| 4 | 2866.91 | 0.000142 | 2.79±0.87 | 1.57±0.36 | 1.78* |
| 5 | 4236.95 | 0.000153 | 1.72±0.44 | 2.57±0.59 | 0.67* |
| 6 | 3365.06 | 0.000158 | 1.78±0.44 | 3.05±0.98 | 0.58* |
| 7 | 1620.19 | 0.000169 | 5.93±2.32 | 2.86±0.92 | 2.07* |
| 8 | 3345.15 | 0.000169 | 1.62±0.37 | 2.29±0.48 | 0.71 |
| 9 | 2664.53 | 0.000229 | 8.73±4.52 | 3.01±1.48 | 2.90* |
| 10 | 3318.21 | 0.000249 | 1.83±0.46 | 3.61±1.48 | 0.51* |
| 11 | 862.58 | 0.000499 | 17.21±11.45 | 42.56±21.88 | 0.40* |
| 12 | 4255.72 | 0.000499 | 1.43±0.19 | 1.83±0.34 | 0.78 |
| 13 | 6637.31 | 0.000721 | 0.73±0.13 | 1.3±0.53 | 0.56* |
| 14 | 846.62 | 0.000749 | 7.08±1.74 | 12.12±4.71 | 0.58* |
| 15 | 878.64 | 0.00253 | 17.46±11.03 | 34.94±17.13 | 0.50* |
| 16 | 2888.06 | 0.00253 | 2.19±0.51 | 1.63±0.43 | 1.34 |
| 17 | 2686.51 | 0.00301 | 2.27±0.49 | 1.74±0.39 | 1.30 |
| 18 | 1074.14 | 0.00448 | 4.46±1.19 | 7.38±3.33 | 0.60* |
| 19 | 1306.7 | 0.00573 | 4.15±0.91 | 5.62±1.64 | 0.74 |
| 20 | 818.31 | 0.00751 | 6.14±1.23 | 7.93±2.04 | 0.77 |
| 21 | 868.05 | 0.0107 | 5.92±1.72 | 9.72±4.89 | 0.61* |
| 22 | 5761.17 | 0.0173 | 0.98±0.26 | 0.75±0.2 | 1.31 |
| 23 | 1549.23 | 0.0256 | 4.55±2.51 | 2.84±0.67 | 1.60* |
| 24 | 2109.3 | 0.0256 | 2.62±0.47 | 2.15±0.58 | 1.22 |
| 25 | 3283.41 | 0.0256 | 1.93±0.32 | 2.28±0.46 | 0.85 |
| 26 | 3238.15 | 0.0274 | 1.78±0.42 | 2.21±0.57 | 0.81 |
| 27 | 1090.3 | 0.0287 | 4.74±2 | 7.01±3.16 | 0.68 |
| 28 | 1262.68 | 0.0289 | 3.99±0.89 | 4.97±1.36 | 0.80 |
| 29 | 1425.85 | 0.0332 | 10.9±3.67 | 8.19±2.76 | 1.33 |
| 30 | 4077.55 | 0.0337 | 1.58±0.38 | 1.3±0.3 | 1.22 |
| 31 | 3978.14 | 0.0348 | 1.62±0.36 | 1.32±0.37 | 1.23 |
| 32 | 1411.31 | 0.0354 | 5.24±1.11 | 4.39±0.99 | 1.19 |
| 33 | 4128.21 | 0.0434 | 1.51±0.38 | 1.26±0.23 | 1.20 |
| 34 | 4650.88 | 0.0437 | 1.34±0.64 | 1.89±0.79 | 0.71 |
| 35 | 6438.84 | 0.0486 | 0.63±0.11 | 0.76±0.23 | 0.83 |

### The difference of potential biomarkers screening between two types of blood collecting tubes

We used ClinProTools to find the different peaks among patient group, control group, patient coagulant group and control coagulant group, and we identified 68 distinguishable peaks of which 56 significantly differed among the groups (P<0.05, table 5). Among this 56 peaks, 20 peaks where the difference between control and patient in coagulant group were reversed when compared with non-coagulant group (Figure 6C), 18 peaks enlarged (Figure 6D) and 18 peaks narrowed (Figure 6F). In addition, in the principal component analysis, the bivariate plot performed only a few overlapping areas between patient group and control group and between patient coagulant group and control coagulant group were found, indicating that the groups were accurately (Figure 6B).

**Table 5.** Peaks showing significant differences in abundance across samples from patient, control, patient coagulant and control coagulant.

| Peak | Mass | P value | patient | control | patient<br>coagulant | control<br>coagulant |
| --- | --- | --- | --- | --- | --- | --- |
| 1 | 1448.12 | < 0.000001 | 5.05±1.03 | 7.98±1.95 | 5.53±1.51 | 8.74±1.75 |
| 2 | 5872.35 | < 0.000001 | 1.61±0.37 | 0.91±0.19 | 1.51±0.45 | 1.5±0.35 |
| 3 | 5761.41 | < 0.000001 | 1.1±0.21 | 0.67±0.09 | 0.98±0.26 | 0.8±0.18 |
| 4 | 1620.27 | < 0.000001 | 5.82±1.7 | 5.5±2.04 | 5.93±2.32 | 2.94±0.88 |
| 5 | 4236.93 | < 0.000001 | 1.8±0.32 | 1.38±0.29 | 1.72±0.45 | 2.58±0.6 |
| 6 | 862.64 | < 0.000001 | 9.32±5.22 | 35.13±23.11 | 17.21±11.45 | 39.84±18.68 |
| 7 | 4215.73 | < 0.000001 | 9.13±3.79 | 3.35±1.65 | 6.08±2.83 | 8.47±3.69 |
| 8 | 5811.58 | < 0.000001 | 0.98±0.22 | 0.64±0.09 | 0.85±0.24 | 0.86±0.16 |
| 9 | 5911.99 | 0.00000192 | 3.85±1.44 | 1.97±0.92 | 4.26±2.22 | 5.21±2.17 |
| 10 | 7773.9 | 0.00000225 | 1.21±0.38 | 1.9±1.02 | 1.03±0.68 | 3.44±1.39 |
| 11 | 3318.28 | 0.00000225 | 1.86±0.4 | 3.23±1.06 | 1.83±0.46 | 3.69±1.47 |
| 12 | 3365.22 | 0.00000266 | 1.52±0.25 | 1.45±0.29 | 1.74±0.46 | 3.09±0.99 |
| 13 | 4970.09 | 0.00000266 | 1.08±0.27 | 1.8±0.58 | 0.89±0.14 | 0.81±0.24 |
| 14 | 2664.63 | 0.00000271 | 7.31±3.06 | 5.49±2.96 | 8.73±4.52 | 3.1±1.46 |
| 15 | 9300.65 | 0.0000029 | 0.81±0.29 | 1.23±0.59 | 0.71±0.4 | 2.23±0.92 |
| 16 | 878.72 | 0.00000382 | 10.37±5.5 | 28±17.8 | 17.48±11.02 | 33.05±15.31 |
| 17 | 2866.92 | 0.0000111 | 3.35±1.81 | 2.02±0.6 | 2.77±0.88 | 1.59±0.37 |
| 18 | 4272.61 | 0.0000132 | 1.55±0.26 | 1.12±0.32 | 1.62±0.5 | 1.92±0.47 |
| 19 | 4255.82 | 0.0000243 | 1.47±0.24 | 1.2±0.28 | 1.44±0.2 | 1.84±0.35 |
| 20 | 846.67 | 0.0000247 | 5.96±1.32 | 11.27±6.43 | 7.05±1.78 | 11.68±4.39 |
| 21 | 1074.19 | 0.0000276 | 3.81±1.12 | 7.26±2.84 | 4.45±1.19 | 6.95±2.8 |
| 22 | 6638.28 | 0.0000366 | 1.02±0.28 | 0.73±0.15 | 0.73±0.13 | 1.32±0.54 |
| 23 | 4987.4 | 0.0000504 | 0.93±0.19 | 1.39±0.41 | 0.87±0.18 | 0.79±0.16 |
| 24 | 4199.33 | 0.0000843 | 2.01±0.54 | 1.24±0.37 | 1.7±0.45 | 1.82±0.44 |
| 25 | 1068.58 | 0.000134 | 4.61±1.11 | 8.15±3.46 | 6.49±3.42 | 7.39±2.78 |
| 26 | 1262.77 | 0.000138 | 3.71±0.68 | 5.05±0.94 | 3.97±0.92 | 4.92±1.38 |
| 27 | 1425.91 | 0.000145 | 13.89±4.01 | 8.88±1.91 | 10.9±3.67 | 8.31±2.78 |
| 28 | 1090.34 | 0.00019 | 4.1±1.04 | 6.88±2.4 | 4.74±2 | 6.68±2.88 |
| 29 | 1306.68 | 0.00019 | 3.84±0.61 | 5.14±1.21 | 4.14±0.93 | 5.51±1.62 |
| 30 | 2109.37 | 0.000285 | 3±0.63 | 2.09±0.61 | 2.62±0.47 | 2.15±0.61 |
| 31 | 3212.5 | 0.00043 | 1.71±0.33 | 1.64±0.36 | 2.04±0.59 | 2.53±0.76 |
| 32 | 2888.22 | 0.000849 | 2.13±0.42 | 1.7±0.33 | 2.17±0.53 | 1.62±0.44 |
| 33 | 2686.49 | 0.00116 | 2.3±0.49 | 2.26±0.63 | 2.32±0.48 | 1.78±0.38 |
| 34 | 4059.41 | 0.00146 | 1.84±0.35 | 1.35±0.32 | 1.53±0.31 | 1.45±0.43 |
| 35 | 1468.94 | 0.00193 | 5.79±1.98 | 7.67±2.9 | 6.62±3.56 | 4.64±1.25 |
| 36 | 867.86 | 0.00193 | 5.3±0.91 | 7.64±3.46 | 5.88±1.68 | 9.5±4.83 |
| 37 | 4077.63 | 0.00199 | 1.56±0.28 | 1.22±0.34 | 1.63±0.34 | 1.32±0.28 |
| 38 | 4127.88 | 0.00219 | 1.58±0.37 | 1.17±0.26 | 1.51±0.38 | 1.31±0.22 |
| 39 | 3957.41 | 0.00232 | $2.56 \pm 0.82$ | $2.07 \pm 0.52$ | $1.86 \pm 0.55$ | $1.59 \pm 0.57$ |
| 40 | 2957.6 | 0.00243 | $2.22 \pm 0.61$ | $1.69 \pm 0.4$ | $2.46 \pm 0.96$ | $2.24 \pm 0.73$ |
| 41 | 810.14 | 0.00453 | $11.95 \pm 2.53$ | $10.25 \pm 2.87$ | $13.07 \pm 4.6$ | $18.76 \pm 9.27$ |
| 42 | 894.82 | 0.00454 | $6.16 \pm 2.03$ | $9.34 \pm 4.71$ | $7.58 \pm 3.84$ | $9.89 \pm 4.01$ |
| 43 | 5343.26 | 0.00454 | $1.21 \pm 0.3$ | $0.89 \pm 0.33$ | $1.31 \pm 0.33$ | $1.21 \pm 0.38$ |
| 44 | 1411.34 | 0.00519 | $5.8 \pm 1.24$ | $4.66 \pm 1$ | $5.24 \pm 1.11$ | $4.44 \pm 0.99$ |
| 45 | 1868.94 | 0.00519 | $2.8 \pm 0.49$ | $3.1 \pm 1.18$ | $2.9 \pm 0.61$ | $2.28 \pm 0.53$ |
| 46 | 6439.19 | 0.00547 | $0.78 \pm 0.17$ | $0.63 \pm 0.13$ | $0.63 \pm 0.1$ | $0.77 \pm 0.23$ |
| 47 | 1084.97 | 0.00635 | $4.17 \pm 1.39$ | $6.14 \pm 1.71$ | $4.99 \pm 2.01$ | $5.5 \pm 2.02$ |
| 48 | 3283.37 | 0.00639 | $2.01 \pm 0.45$ | $1.77 \pm 0.31$ | $1.93 \pm 0.32$ | $2.28 \pm 0.47$ |
| 49 | 818.35 | 0.0124 | $6.77 \pm 2.23$ | $6.01 \pm 1.47$ | $6.09 \pm 1.25$ | $7.98 \pm 2.08$ |
| 50 | 1052.27 | 0.0132 | $4.27 \pm 0.97$ | $6.27 \pm 2.52$ | $4.73 \pm 1.28$ | $5.52 \pm 2.08$ |
| 51 | 1549.33 | 0.0148 | $4.72 \pm 2.57$ | $3.45 \pm 0.98$ | $4.63 \pm 2.44$ | $3.05 \pm 0.84$ |
| 52 | 1279.83 | 0.0172 | $4.14 \pm 0.78$ | $5.47 \pm 1.83$ | $4.62 \pm 1.45$ | $5.09 \pm 1.35$ |
| 53 | 826.13 | 0.0197 | $9.45 \pm 3.73$ | $6.71 \pm 1.54$ | $7.25 \pm 1.41$ | $8.72 \pm 3.41$ |
| 54 | 4096.5 | 0.0251 | $3.42 \pm 1.28$ | $2.21 \pm 1.03$ | $2.87 \pm 1.06$ | $2.49 \pm 0.86$ |
| 55 | 835.34 | 0.0309 | $7.06 \pm 1.36$ | $6.27 \pm 1.61$ | $6.47 \pm 2.12$ | $8.55 \pm 2.71$ |
| 56 | 884.2 | 0.0415 | $6.73 \pm 3.18$ | $8.33 \pm 3.69$ | $8.81 \pm 5.73$ | $11.19 \pm 5.49$ |

**Figure 6.**
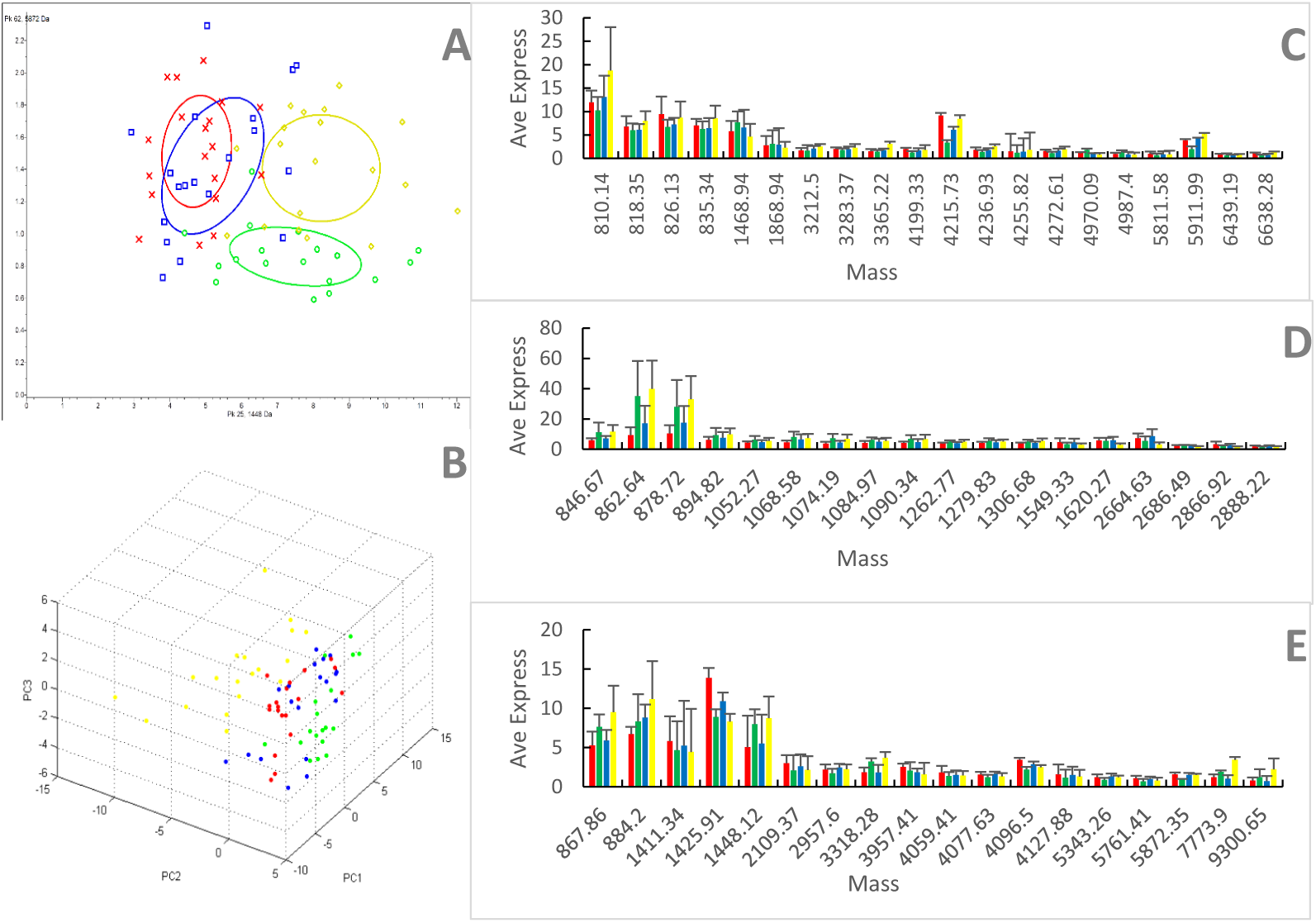
Comparative profiling of serum peptides among four different groups. Bivariate plot (A) and 3D plot (B) of patient group (red), control group (green), patient coagulant group (blue) and control coagulant group (yellow) in the principal component analysis. In coagulant group, the difference between control and patient was reversed (C), enlarged (D), and narrowed (E) compared with non-coagulant group.

## Discussion

The emerging technology, proteomics, is an effective high-throughput research mode, which is able to investigate the overall protein expression changes of the class and quantity dynamically and quantitatively, and to obtain some new protein markers as well as new key molecular^26,27^. It greatly enriches the optimal selections and combinations of diagnostic markers, which provides new ideas for us to seek specific diagnostic markers^28^, study the pathogenesis of the complex disease comprehensively, test the prognosis and recurrence of disease, and screen to identify the therapeutic targets^29^. Therefore, it’s significant to find out the most representative protein marker. In the present study, we selected blood samples from different blood diseases (including multiple myelomatosis, non-Hodgkin lymphoma, and leukemia), different genders and different ages (from 57-73) to expand the apply scope of the experiment. In addition, we detected in normal people as control subjects to make horizontal and vertical comparisons. The result showed that 26 peaks were screened out when we collected the samples with non-additive tubes, and some of them can be potential markers. They are 5871.42, 4215.81, 1448.28, 1425.99, 5912.25, 3318.58, 4970.76, 1074.14, 1090.44, 4988.5, 862.72, 1068.65, 878.84, 846.78, 4096.62, 9301.2, 2866.95, 7773.61, and 894.94. When we collected with coagulant tubes, 35 peaks were screened out between the disease and normal groups. But the peak 5871.42 did not showed up, and some of the peaks (such as 4215.91, 1425.99, and 5912.25) had little difference between the different groups. It means that the detected protein expression value was significantly changed after the addition of coagulant for pre-treatment in both the disease group and normal group. And there were significant difference among the high peaks found. It told us that adding coagulant would obviously influence the peaks found by MS, and influence the further selection of the significant markers, which would have a negative impact on the clinical diagnosis. Moreover, related studies have shown that the detection was effected by the centrifugation time, freshness of the samples^30^, types of the blood collection tubes^23^, delays in centrifugation as well^31^. As we can see, processing affects the detection greatly during the blood sample collection. However, there is no standardization of the procedure^32^. Some researchers use the non-additive tubes, the others choose to use coagulant activator tubes. Although in 2005, HUPO has come up with a standardized blood-collecting plan^33^, the selection of the type of the tubes has not been clearly ruled. That was a long time ago, so that it’s necessary and urgent to put forward a new standard. What’s more, we also found that the addition of coagulant had more effect on the MS result in the normal group while had less effect in the patient group. This may be related to the wide variation of serum proteins in normal blood, while the serum proteins in patients are common due to the co-occurrence of certain diseases. Since the addition of coagulant has a great impact on the serum of normal people, it remains to be further studied whether the blood collection method of adding coagulant, which is widely used in clinical practice, is reasonable.

